# LC3B phosphorylation regulates FYCO1 binding and directional transport of autophagosomes

**DOI:** 10.1101/2020.05.15.081638

**Authors:** Jose L. Nieto-Torres, Sean-Luc Shanahan, Romain Chassefeyre, Sara Landeras-Bueno, Sandra E. Encalada, Malene Hansen

## Abstract

Macroautophagy (hereafter referred to as autophagy) is a conserved process that promotes cellular homeostasis through the degradation of cytosolic components, also known as cargo. During autophagy, cargo is sequestered into double-membrane vesicles called autophagosomes, which are predominantly transported in the retrograde direction to the perinuclear region to fuse with lysosomes, thus ensuring cargo degradation [1]. The mechanisms regulating directional autophagosomal transport remain unclear. The ATG8 family of proteins associate with autophagosome membranes [2] and play key roles in autophagy, such as the movement of autophagosomes. This is achieved via the interaction of ATG8 with adaptor proteins, including FYCO1, a protein involved in the anterograde transport of autophagosomes toward the cell periphery [1,3–5]. We previously reported that phosphorylation of LC3B/ATG8 on threonine 50 (LC3B-T50) by the Hippo kinase STK4 is required for autophagy through unknown mechanisms [6]. Here, we show that LC3B-T50 phosphorylation decreases the interaction between LC3B and FYCO1, which in turn regulates the starvation-induced perinuclear positioning of autophagosomes. Moreover, non-phosphorylatable LC3B-T50A aberrantly switches the predominant retrograde movement of autophagosomes to anterograde movement towards the cell periphery in multiple cell types, including in mouse primary hippocampal neurons. Our data support a role of a nutrient-sensitive STK4–LC3B–FYCO1 axis in the regulation of the directional transport of autophagosomes via the post-translational regulation of LC3B. Given that autophagy is impaired in many human conditions, including neurodegenerative diseases, our findings may highlight new principles of vesicle transport regulation critical for disease etiology.

**Graphical abstract:** 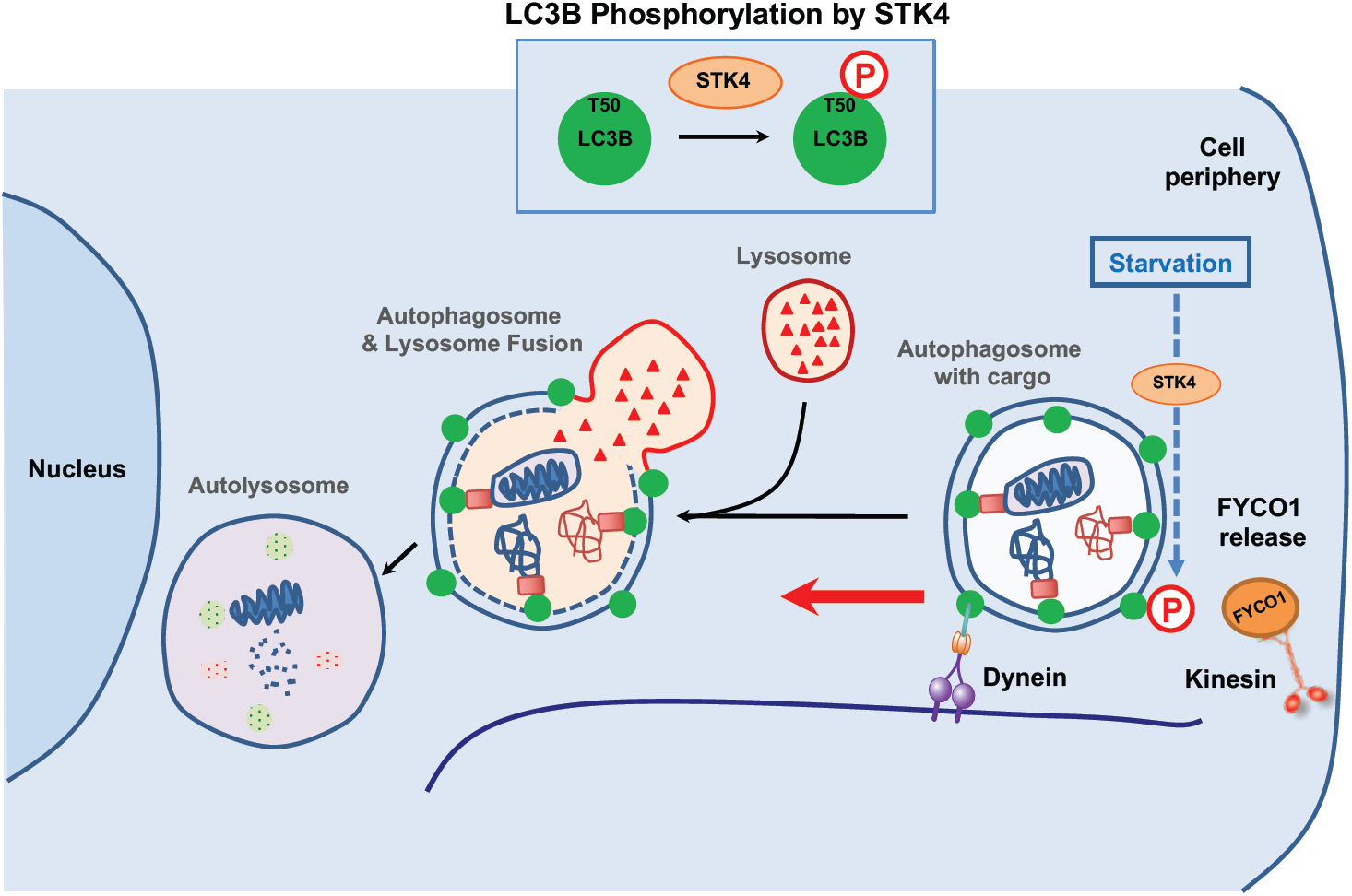

## RESULTS

### LC3B-T50 Phosphorylation Regulates the Interaction Between LC3B and FYCO1

We recently reported that phosphorylation of LC3B-T50 by STK4 is required for autophagy [6]. Specifically, depletion of *Stk4* or expression of a non-phosphorylatable LC3B-T50A mutant causes accumulation of autophagosomes throughout the cell and clustering of lysosomes around the nucleus, leading to a block in autophagy in mouse embryonic fibroblasts (MEFs) and C2C12 mouse myoblasts [6]. We hypothesized that phosphorylation of LC3B might modulate its interaction with binding partners such as autophagy adaptor proteins. To test this, we transiently expressed HA-tagged wild-type (WT) LC3B or the phospho-mutant forms LC3B-T50A (T50A, phospho-deficient) and LC3B-T50E (T50E, phospho-mimetic) in human 293T cells, affinity purified the HA-tagged LC3B proteins, and identified the associated proteins by mass spectrometry (**Figure 1A**). Of the LC3B interactors identified in this analysis, FYCO1, a known autophagy adaptor protein [3,4], showed the most LC3B-T50 phosphorylation-sensitive binding; specifically, FYCO1 binding was ∼2-fold higher to the T50A mutant and ∼5-fold lower to the T50E mutant compared with binding to LC3B-WT (**Figure 1B**).

**Figure 1.**
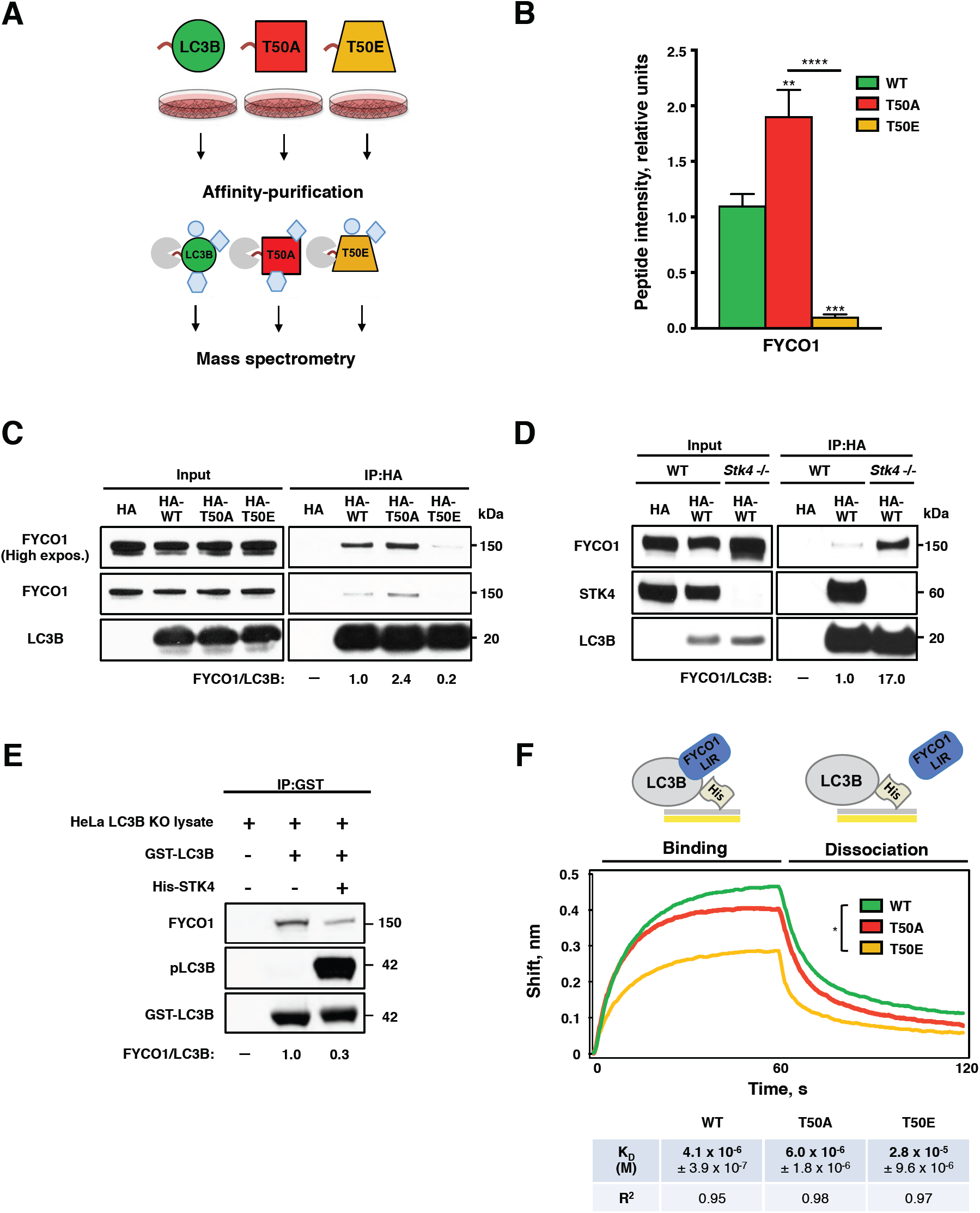
LC3B phosphorylation modulates FYCO1–LC3B interactions. (**A**) Schematic of pull-down assays performed in HEK 293T cells expressing WT (wild-type), T50A (phospho-deficient mutant), and T50E (phospho-mimetic mutant) LC3B proteins to identify LC3B phosphorylation-dependent interactors by mass spectrometry. (**B**) Quantification of FYCO1 binding to LC3B proteins determined by mass spectrometry. Mean ± SD of three technical replicates. **p<0.0021, ***p<0.0002, ****p<0.0001 by one-way ANOVA. (**C**) Representative western blot of FYCO1 co-immunoprecipitated with HA-tagged LC3B proteins expressed in HeLa LC3B-KO cells (n=3). (**D**) Representative western blot of FYCO1 co-immunoprecipitated with HA-tagged LC3B proteins in STK4-deficient and WT mouse embryonic fibroblasts (n=3). **(E)** Representative western blot of FYCO1 co-immunoprecipitated with non-phosphorylated or *in vitro-*phosphorylated GST-LC3B (n=4). GST-LC3B panel shows Coomassie staining. (**C-E**) Values below blots are protein ratios in relative units. (**F**) Biolayer interferometry assay of LC3B proteins binding to a peptide containing FYCO1-LIR and adjacent amino acids [4]. Mean K_D_ ± SEM of three technical replicates and R^2^ for binding curve fit are shown. *p<0.05 by one-way ANOVA.

To validate these results, we generated an LC3B knockout (KO) of HeLa cells using CRISPR-CAS9 technology (**Figure S1A**), and expressed LC3B-WT, T50A, or T50E in these LC3B-KO cells. The autophagy-related phenotypes of the LC3B phospho-mutants in the LC3B-KO cells were similar to those reported for mouse cells [6], including an increase in autophagosomes numbers in LC3B KO cells expressing T50A that was insensitive to the autophagy inhibitor Bafilomycin A (**Figure S1B-D**). We therefore used the HeLa LC3B-KO cell line throughout the rest of the study, unless otherwise indicated. Immunoprecipitation of LC3B (**Figure 1C**) or FYCO1 (**Figure S2A**) followed by western blotting for the reciprocal protein confirmed the differential interaction of FYCO1 with LC3B-WT and T50A. Likewise, depletion of *Stk4* in MEFs caused a striking increase in FYCO1–LC3B binding (**Figure 1D**). Finally, *in vitro* phosphorylation of LC3B on T50, verified with a phosphorylation-specific antibody [6], reduced LC3B interaction with FYCO1 compared with non-phosphorylated LC3B (**Figure 1E**). Collectively, these data indicate that STK4-mediated phosphorylation of LC3B decreases its interaction with FYCO1, suggesting a role for FYCO1 in STK4–LC3B-mediated autophagy regulation.

LC3B is known to directly interact with FYCO1 through its LC3-interacting region (LIR) [3,4]. Within the LC3B–FYCO1 binding interface, amino acid T50 of LC3B is located in close proximity to two aspartic acid residues of FYCO1 (**Figure S2B**). We therefore speculated that LC3B-T50 phosphorylation may directly affect FYCO1–LC3B binding. To test this, we performed biolayer interferometry to measure the binding affinities between LC3B-WT or phospho-mutants (bacterially expressed and purified) and a FYCO1 33 amino-acid peptide (1265–1298) spanning the LIR domain and its adjacent aspartic acids [4]. The binding affinity of LC3B-WT and the FYCO1 peptide was in the micromolar range (K_D_ ∼4.1 × 10^−6^ M) (**Figure 1F**), similar to previous reports [7]. In contrast, the binding affinity of the phospho-mimetic T50E and the FYCO1 peptide was significantly lower (K_D_ ∼2.8 x 10^−5^ M) than that of LC3B-WT (**Figure 1F**), supporting the notion that T50 phosphorylation may play a key role in blocking the direct interaction between LC3B and the FYCO1-LIR region. T50A bound to the FYCO1 peptide with an affinity (K_D_ ∼6.0 x 10^−6^ M) in the same micromolar range as LC3B-WT (**Figure 1F**), which was expected because Hippo kinases are not expressed in bacteria [8] and LC3B-WT and T50A would thus behave in a similar manner. Collectively, these data suggest that phosphorylation of LC3B-T50 directly decreases LC3B binding affinity for FYCO1. Overall, these results indicate that phosphorylation of LC3B regulates the interaction between LC3B and FYCO1 and decreases LC3B-FYCO1 binding affinity.

### Subcellular Localization of Autophagosomes is Regulated by LC3B-T50 Phosphorylation in a FYCO1-Dependent Manner

As FYCO1 associates with LC3B on autophagosomes to regulate their transport [3,4], we next asked if LC3B-T50 phosphorylation status would affect FYCO1 localization to autophagosomes within cells. We counted LC3B-positive punctae as representative of autophagosomes, since lipidation-deficient versions of WT and phospho-mutant LC3Bs, which cannot associate with autophagosomes membranes [2], were diffusely localized (**Figure S2C**). These lipidation-deficient versions of WT and phospho-mutant LC3Bs maintained their differential binding to FYCO1 (**Figure S2D**), suggesting that LC3B lipidation may not be essential for LC3B phosphorylation to regulate LC3B-FYCO1 binding. Notably, the percentage of LC3B/FYCO1 double-positive punctae relative to total LC3B punctae was higher in T50A, and lower in T50E, as compared to those in LC3B-WT-expressing cells (**Figure 2A-B and S2E**), indicating that LC3B-T50 phosphorylation decreases whereas de-phosphorylation increases the association of FYCO1 with autophagosomes, consistent with the LC3B-FYCO1 interaction and binding-affinity data (**Figure 1**).

**Figure 2.**
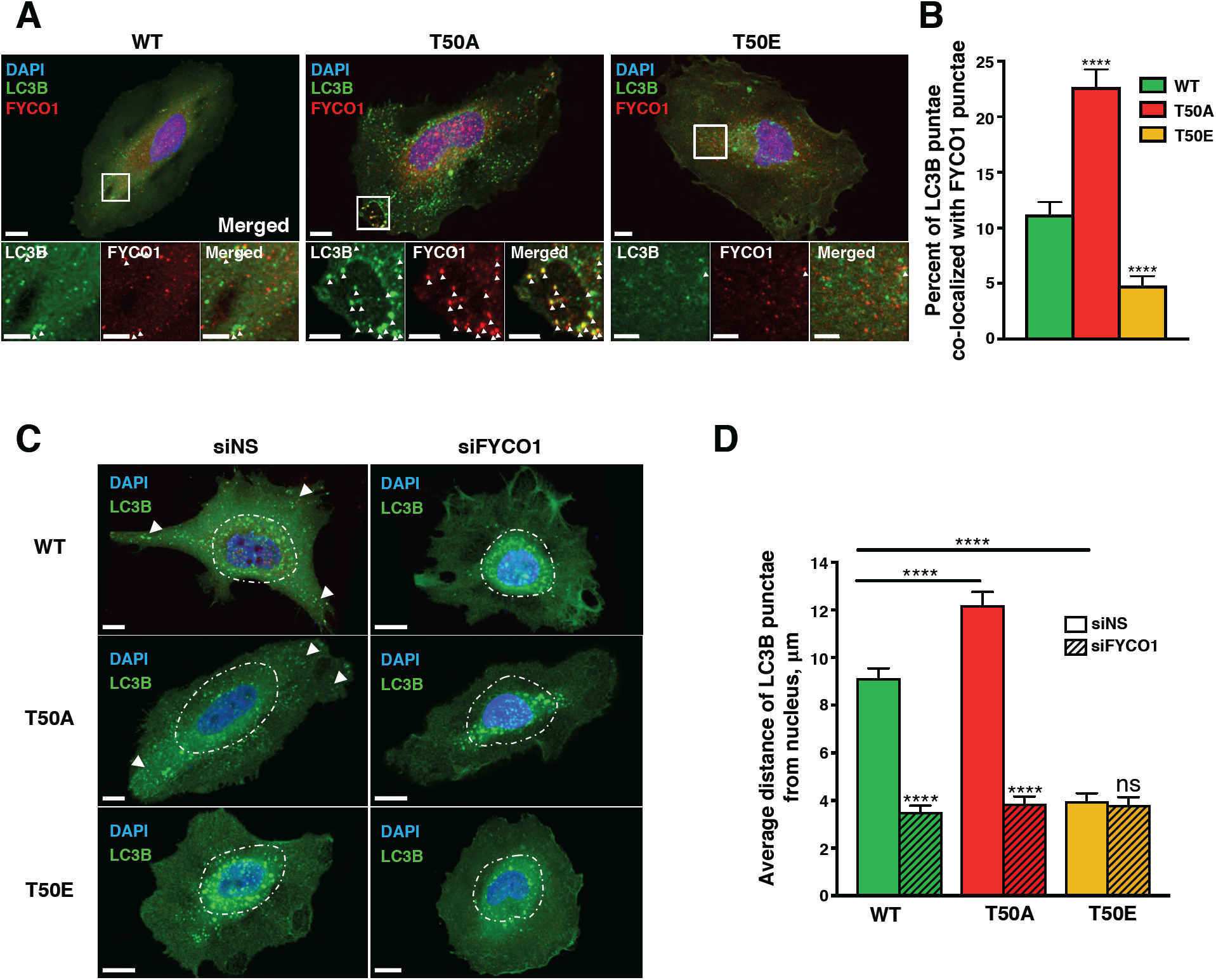
Subcellular localization of LC3B phospho-mutants is differentially regulated by FYCO1. **(A)** Representative immunofluorescence micrographs of HeLa-KO cells expressing HA-LC3B-WT, T50A, or T50E co-stained for FYCO1. White arrowheads indicate co-localized LC3B and FYCO1 punctae. **(B)** Percentage of LC3B punctae co-localizing with FYCO1. Mean ± SEM, ****p<0.0001 by two-way ANOVA (n=16–27 cells from three experiments). **(C)** Representative immunofluorescence micrographs of LC3B-WT, T50A, or T50E punctae in control (siNS) or siFYCO1-treated HeLa cells. Dashed line defines 10 μm from the outer nuclear edge. Scale bar = 10 μm. White arrowheads point to LC3B-positive punctae in peripheral locations of the cell. (**D**) Average distance between LC3B-positive punctae and the nucleus. Mean ± SEM of n=20– 34 cells from two experiments. ****p<0.0001 by two-way ANOVA.

As FYCO1 regulates anterograde transport of autophagosomes [3], we next asked whether the LC3B phosphorylation-dependent association with FYCO1 affected the subcellular positioning of autophagosomes. Immunofluorescence microscopy and imaging analysis revealed that the average distance between LC3B-positive punctae and the nucleus was significantly shorter in T50E-expressing cells (∼4.0 μm), than in cells expressing LC3B-WT (∼9.2 μm), whereas T50A showed a longer average distance (∼12.2 μm). Notably, the dispersal of LC3B punctae to the cell periphery in WT and T50A-expressing cells was FYCO1-dependent, as siRNA-mediated FYCO1 knockdown (**Figure S2F**) resulted in perinuclear accumulation of LC3B punctae (**Figure 2C-D and S3A**). These results show that the subcellular localization of LC3B punctae is regulated by LC3B phosphorylation in a FYCO1-dependent manner.

### Starvation-Induced Perinuclear Clustering of Autophagosomes is Regulated Via a STK4– LC3B–FYCO1 Axis

Starvation is a robust inducer of autophagy and is known to enhance perinuclear positioning of autophagosomes [9]. We next investigated the effects of starvation on STK4–LC3B–FYCO1-mediated control of autophagosomal localization. Combined serum and amino-acid deprivation of HeLa cells resulted in a ∼3-fold increase in the levels of phosphorylated (active) STK4 (STK4-pT183) [10] (**Figure 3A**), consistent with previous findings that the Hippo pathway is activated following serum deprivation [11]. In turn, siRNA-mediated depletion of STK4 (**Figure S3B**) prevented starvation-induced perinuclear clustering of LC3B-positive punctae in LC3B-WT– expressing cells (**Figure 3B-C and S3C**), suggesting that STK4 activation is required for the starvation-induced perinuclear positioning of autophagosomes. Consistent with these results, starvation did not change the dispersed or perinuclear subcellular localization of LC3B-positive punctae in T50A or T50E cells, respectively (**Figure 3D-E and S3D**). Collectively, these results indicate that starvation induces STK4-mediated LC3B phosphorylation-dependent positioning of autophagosomes in mammalian cells.

**Figure 3.**
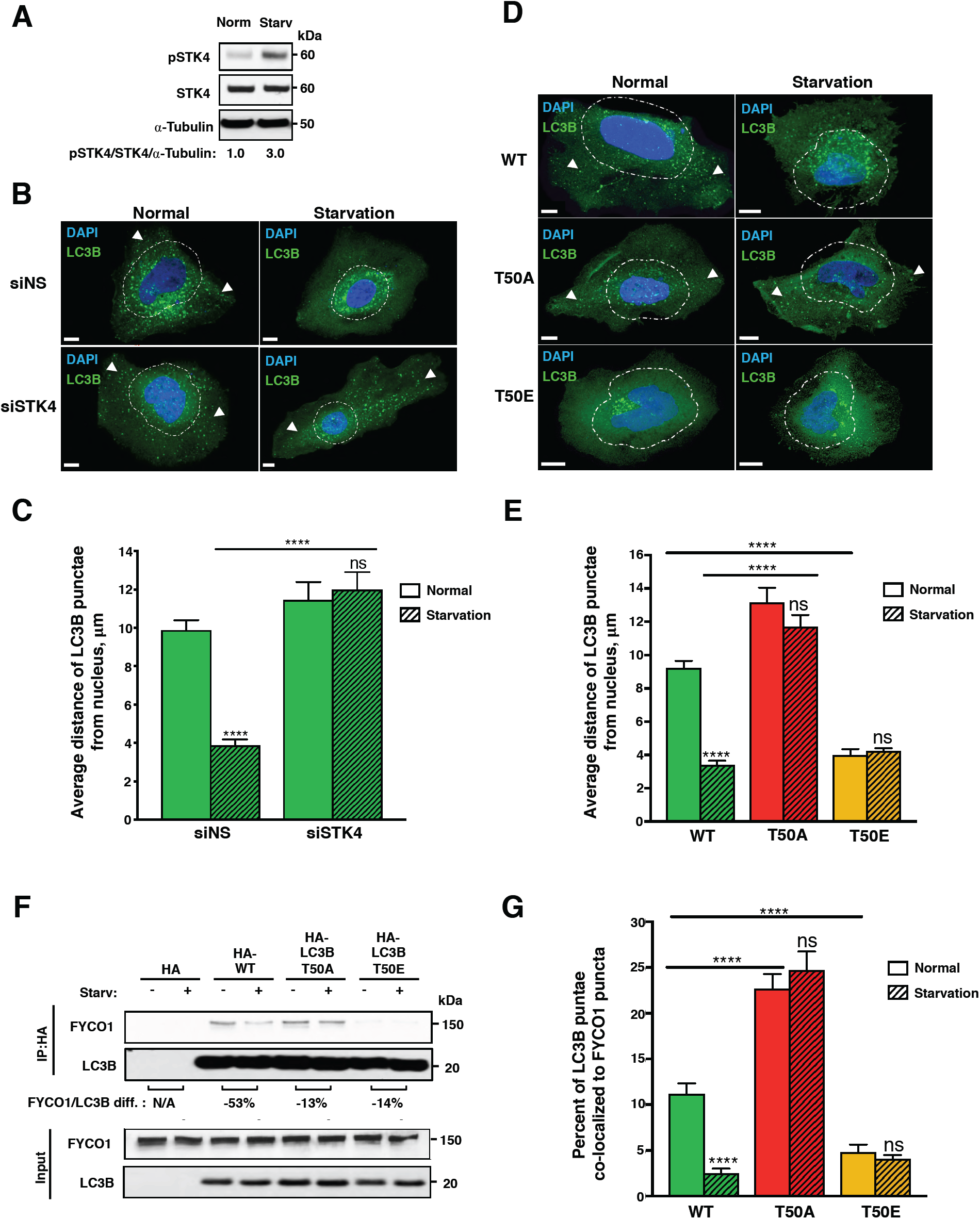
Starvation induces perinuclear localization of LC3B punctae in an LC3B phosphorylation- and FYCO1-dependent manner. (**A**) Representative western blot of total and phosphorylated STK4-T183 in HeLa LC3B-KO cells incubated in normal or starvation medium (n=3). Values indicate pSTK4:STK4 protein ratio normalized to α-tubulin. (**B**) Representative immunofluorescence micrographs of LC3B punctae in HeLa LC3B-KO cells expressing control or STK4-targeting siRNA before or 1 h after incubation in starvation medium. Scale bar = 10 μm. White arrowheads point to LC3B-positive punctae in peripheral locations of the cell. (**C**) Quantification of distance between LC3B punctae and the nucleus. Mean ± SEM of n=24–34 cells from three experiments. ***p<0.0002 by two-way ANOVA. (**D and E**) Representative immunofluorescence micrographs (**D**) and quantification of distance from the nucleus (**E**) of LC3B punctae in HeLa cells expressing the indicated LC3B proteins. Scale bar = 10 μm. White arrowheads point to LC3B-positive punctae in peripheral locations of the cell. Mean ± SEM of n=18–23 cells from two experiments. ****p<0.0001 by two-way ANOVA. (**F**) Representative western blot of FYCO1 co-immunoprecipitated with HA-tagged LC3B proteins expressed in HeLa LC3B-KO cells before and 1 h after incubation in normal or starvation medium. Values under blots indicate percent decrease in LC3B:FYCO1 protein ratio after starvation (n=2). (**G**) Percentage of LC3B punctae co-localizing with FYCO1 in HeLa LC3B-KO cells before or 1 h after incubation in starvation medium. Mean ± SEM of n=16–27 cells from two experiments. ****p<0.0001 by two-way ANOVA.

To determine whether starvation affects LC3B–FYCO1 binding, we performed co-immunoprecipitation experiments and immunofluorescence microscopy of cells expressing LC3B-WT or phospho-mutants grown under normal or starvation conditions. In LC3B-WT– expressing cells, starvation caused a 53% reduction in FYCO1 co-immunoprecipitation with LC3B (**Figure 3F**) and significantly decreased the percentage of LC3B-positive punctae co-localizing with FYCO1 (**Figure 3G and S3E**) In contrast, starvation of cells expressing T50A or T50E only slightly reduced (13-14%) the interaction between FYCO1 and either LC3B phospho-mutant (**Figure 3F**) and had no effect on the co-localization of FYCO1 with LC3B phospho-mutant–positive punctae (**Figure 3G and S3E**). Thus, starvation decreased the association between FYCO1 and LC3B in an LC3B-phosphorylation dependent manner. Overall, these data indicate that nutrient deprivation regulates the STK4–LC3B–FYCO1 axis to control the subcellular positioning of autophagosomes.

### LC3B Phosphorylation Regulates the Directional Transport of Autophagosomes in Mammalian Cells and Neurons

Previous work indicated that FYCO1 promotes the anterograde transport of autophagosomes via its interactions with LC3B and the motor protein kinesin [3,12]. Since we showed above that LC3B-T50 phosphorylation regulated both the binding of LC3B to FYCO1 as well as autophagosome positioning, we next asked whether LC3B-T50 phosphorylation plays a role in autophagosomal transport. To investigate this, we used super-resolution (SR) live-cell imaging to characterize the intracellular transport of LC3B-positive vesicles in HeLa LC3B KO cells expressing GFP-tagged LC3B proteins (**Figure 4A**). Consistent with previous reports [13,14], GFP-LC3B WT vesicles moved primarily in the retrograde direction towards the perinuclear region, as was the case for GFP-T50E particles (**Figure 4B and Video S1 and S3**). Notably, T50E vesicles also displayed significant switches in directionality not exhibited by either GFP-LC3B WT or GFP-T50A vesicles, suggesting impaired coordination between anterograde and retrograde motors associated to those vesicles (**Figure S4A and Video S3**). In contrast, GFP-T50A vesicles were mainly transported in an anterograde direction towards the cell periphery, suggesting that phosphorylation of LC3B is important for perinuclear positioning (**Figure 4B and Video S2**). Overall, these results indicate that LC3B phosphorylation regulates coordination of directional transport of autophagosomes in HeLa cells and is required for retrograde transport.

**Figure 4.**
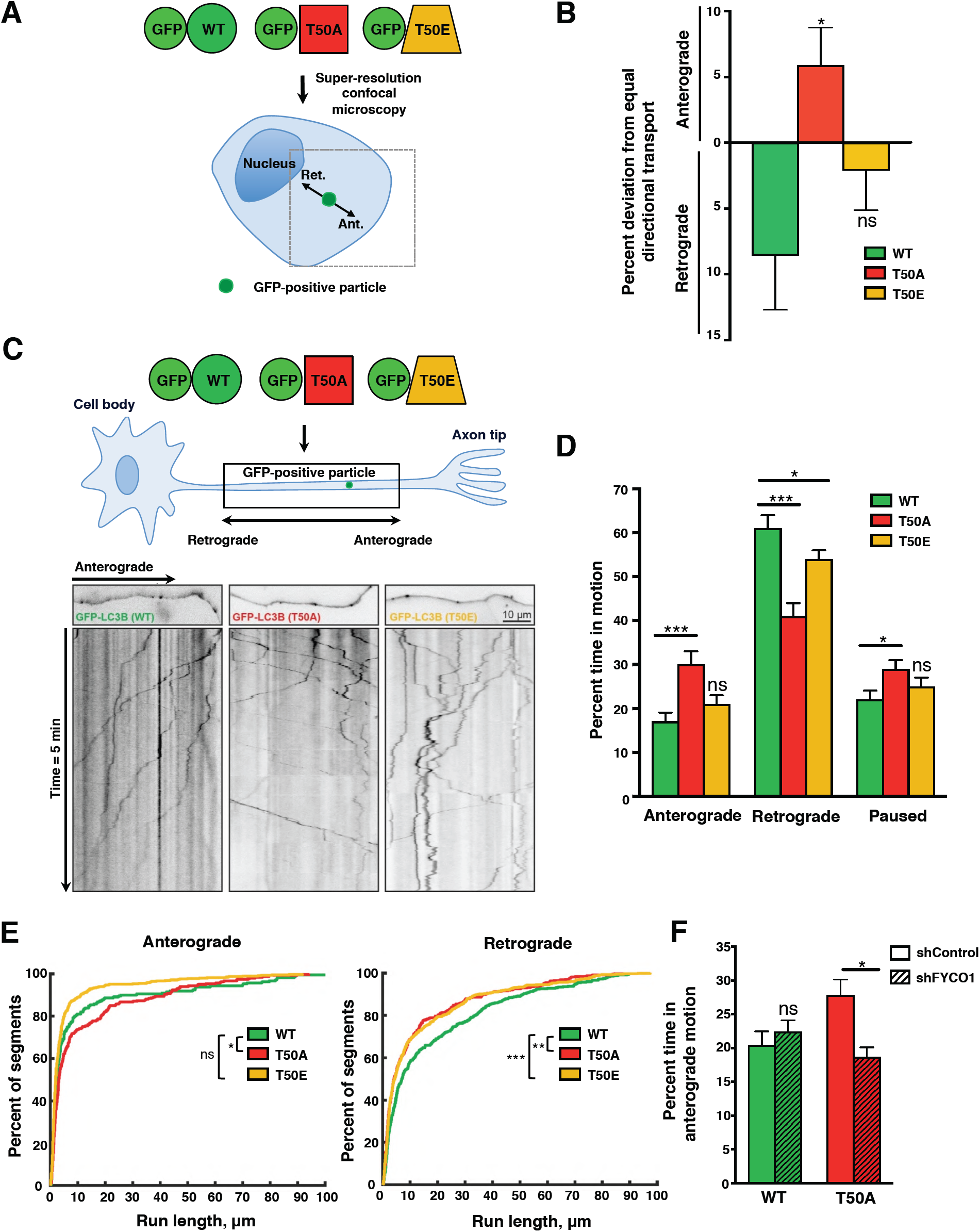
Directional transport of LC3B punctae is regulated by the LC3B phosphorylation. (**A**) Schematic illustration of experimental setup for HeLa LC3B-KO cells. Directional anterograde (Ant) or retrograde (Ret) movement of LC3B-positive vesicles is depicted. (**B**) Net anterograde or retrograde movement of LC3B-positive vesicles per cell determined by SR imaging. Mean ± SEM of n=9–10 cells from two experiments. Percent deviation from equal directional transport was calculated by subtracting the percentage of anterograde-moving from the retrograde-moving LC3B-positive vesicles. Absolute values for net directional transport are shown. *p<0.0332 by one-way ANOVA. (**C**) Schematic illustration of experimental setup for primary mouse hippocampal neurons. Lower images show representative first frame pictures and kymographs of the indicated GFP-positive vesicles (pseudo-black lines). (**D**) Quantification of directional transport of LC3B-positive vesicles in mouse primary neurons. Data are presented as the mean percent time spent stationary (pause), anterograde or retrograde motion for each LC3B-positive particle. Mean ± SEM of n=26–29 cells from two experiments. ***p<0.001 by non-parametric permutation Student’s t-test. (**E**) Cumulative frequency graphs of anterograde and retrograde run lengths of GFP-LC3B-positive vesicles in mouse primary neurons. n=26–29 cells from two experiments. *p<0.05, **p<0.01, ***p<0.001 by Kolmogorov–Smirnov test adjusted with Bonferroni. Similar statistically significant comparisons were obtained when analyzing the data by Rank Sum test (data not shown). (**F**) Proportion of time spent in anterograde transport of GFP-LC3B-positive vesicles in mouse primary neurons with or without *Fyco1* knockdown. Mean ± SEM of n=19–23 cells from two experiments. *p<0.05 by non-parametric permutation Student’s t-test.

To test whether the requirement for LC3B phosphorylation to drive the retrograde transport of LC3B-positive vesicles is conserved, we characterized the dynamics of GFP-LC3B vesicles in dissociated cultured mouse hippocampal neurons. Axonal transport is accomplished by the binding and translocation along uniformly polarized microtubules, of kinesin and dynein that move cargoes in anterograde (toward the synapse) and retrograde (toward the soma) directions, respectively [15,16]. Previous studies showed that autophagosomes form in distal regions of axons, and are transported retrogradely toward the soma, where they fuse with lysosomes to drive degradation of various cargoes [17–19]. To test whether LC3B phosphorylation also affected the transport of autophagosomes in primary neurons, we characterized the dynamics of these vesicles in mammalian axons using live pseudo-total internal reflection microscopy (pTIRFM). Consistent with previous reports [17,18], GFP-LC3B WT vesicles moved primarily in the retrograde direction, spending ∼60% of time moving toward the soma (**Figure 4C-D and Video S4**). In contrast, GFP-T50A vesicles spent significantly less time (∼40%) moving in the retrograde direction, and instead spent longer periods moving anterogradely toward the synapse (**Figure 4C-D and Video S5**). Enhanced GFP-T50A vesicle anterograde movement was also observed following quantitative analysis of run lengths (**Figure 4E**), which indicated that the majority (>80%) of GFP-T50A vesicles moved for longer distances in the anterograde direction, and for shorter distances and slower in the retrograde direction, compared to GFP-WT particles (**Figure 4E and S4B**). In contrast, GFP-T50E vesicles moved primarily in a retrograde fashion (**Figure 4C-D and Video S6**), similarly to GFP-LC3B WT vesicles, but with modified dynamics, including moving shorter distances (**Figure 4E**), suggesting that modulation of the state of LC3B phosphorylation can alter the processive movement of these vesicles. The reduced processivity of GFP-T50E vesicles could be either the result or a readout of the higher frequency of directionality reversals (switches) that these vesicles undertake as they moved along axons (**Figure S4C and Video S6**), similar to the increased numbers of switches observed for GFP-T50E vesicles in HeLa LC3B-KO cells (**Figure S4A and Video S3**). Overall, these results indicate that LC3B phosphorylation regulates directional transport of autophagosomes in a conserved manner in axons.

Previous work showed that FYCO1 drives the anterograde movement of LC3B-positive vesicles [3]. Thus, to characterize the mechanistic bases of LC3B-T50 phosphorylation-dependent regulation of autophagosomal transport, we next tested whether anterograde movement of autophagosomes in axons was dependent of FYCO1. Down-regulation of *Fyco1* using shRNAs rescued the increased time that GFP-T50A vesicles spent moving in the anterograde direction (**Figure 4F**). Thus, the LC3B phosphorylation-dependent changes in GFP-LC3B-positive particle anterograde transport in axons require FYCO1. Altogether, our data show that robust retrograde transport of LC3B-positive autophagosomes via FYCO1 binding regulation requires LC3B-T50 phosphorylation, and the absence of this post-translational modification leads to a reversal to-and enhancement of anterograde transport in HeLa cells and in mammalian neurons.

## DISCUSSION

In this study, we demonstrated that STK4-mediated phosphorylation of the autophagy protein LC3B is critical for regulating its interaction with the adaptor protein FYCO1 and for the control of directional autophagosomal transport in mammalian cells. We also showed that nutrient deprivation activates the STK4–LC3B–FYCO1 regulatory axis. These findings expand our fundamental understanding of the molecular regulation of autophagy with possible implications for the numerous biological processes that autophagy has been linked to [20].

We found that STK4-mediated phosphorylation of LC3B at T50 reduced binding of FYCO1 to LC3B, consistent with a recent report [21], as well as FYCO1 localization to autophagosomes. Compatible with the role of FYCO1 in anterograde transport of intracellular vesicles including autophagosomes [3,4,12], we showed that lack of LC3B-T50 phosphorylation resulted in enhanced anterograde transport and aberrant positioning to the cell periphery of LC3B-positive vesicles in both HeLa LC3B KO cells and primary neurons. Given that prevention of LC3B-T50 phosphorylation also increased lysosomal clustering in HeLa LC3B KO cells (**Figure S1D**) [6], these observations could account for the autophagy block observed in LC3B phosphorylation-deficient cells [6]. Based on these results, we speculate that formation of a LC3B-FYCO1-kinesin complex may promote the anterograde and inhibit the retrograde transport of progressing autophagosomes and by extension their premature fusion with lysosomes [22] that migrate to the nuclear periphery [23]. In turn, we propose that, following autophagosome formation, STK4-mediated phosphorylation of LC3B induces FYCO1 dissociation from mature autophagosomes to allow their retrograde transport and ensure lysosomal fusion and cargo degradation. While other adaptor proteins facilitate the processive retrograde transport of autophagosomes [5,24], it remains to be investigated whether LC3B phosphorylation could similarly affect the binding of retrograde transport-promoting adaptors.

While our studies show that phosphorylation of LC3B was critical for driving retrograde transport in LC3B T50A particles, T50E-positive vesicles revealed an unusually high frequency of directional switches in HeLa LC3B KO cells and in primary neurons. It is possible that the T50E mutation, that mimics a permanent phosphorylation state and decreases the affinity of LC3B for FYCO1, may directly impair the tight coordination between kinesin and dynein [25,26], compromising both anterograde and retrograde motility and thus resulting in frequent reversals. Alternatively, a recent report suggested that the T50E mutation may compromise LC3B lipidation [21], which may result in a reduced association of LC3B with autophagosomes, decreasing FYCO1 association and leading to a loss of anterograde-retrograde coordination. Additional experiments will be critical to explain the directional switches produced by LC3B-T50E, as well as to identify potential LC3B-T50 phosphatase(s) involved in autophagosomal trafficking and recruitment of essential transport machinery. Noting that another ATG8 member, LC3A, also interacts with FYCO1 [4], it will also be of interest to investigate if different ATG8 proteins bind FYCO1 in a phosphorylation-dependent manner and participate in directional transport of autophagosomes.

We observed that starvation-induced perinuclear positioning of autophagosomes was dependent on STK4-mediated LC3B phosphorylation. We propose that the autophagosome repositioning process is initiated by starvation-induced activation of STK4, followed by LC3B phosphorylation, dissociation of FYCO1, and inhibition of anterograde transport and/or promotion of retrograde transport. In support of this model, anterograde transport of autophagosomes has been shown to decrease in HeLa cells during starvation [27]. In low-nutrient conditions, activation of STK4 may concomitantly reduce cell proliferation [28] and increase autophagy in an effort to re-establish homeostasis and restore nutrient availability. To this end, the nutrient-sensing complex mTORC2 has recently been identified as a direct inhibitor of STK4 activity [29] and it will be appealing to investigate its potential role as an upstream regulator of the STK4–LC3B–FYCO1 axis. Indeed, it will be of interest to explore the role of this STK4–LC3B–FYCO1 regulatory axis in different biological processes, including in the decline in autophagy observed during aging and in many disease states [9,30,31], where deregulation of autophagosome transport could be a crucial factor.

## ACKNOWLEDGEMENTS

We thank Lee R. Lee and Sviatlana Zaretski for assistance with the affinity-purification experiments, Dr. Caroline Kumsta and Tai Chaiamarit for help with data analysis and figure preparation, members of the Hansen and Encalada labs for helpful discussions and/or critical reading of the manuscript, Drs. Tony Hunter and Jill Meisenhelder (Salk Institute) for helpful discussions, Dr. Alex Rosa Campos (Sanford Burnham Prebys Proteomics Core) for support with the mass-spectrometry analysis, Leslie Boyd (Sanford Burnham Prebys Cell Imaging Core) and Salk Institute Biophotonics Core personnel for help with image acquisition and analysis, and Dr. Erica Ollmann-Saphire (The Scripps Research Institute) for help with the binding affinity assays. J.N.T. was supported by a Fundacion Ramon Areces Postdoctoral Fellowship, and R.C. was supported by the George E. Hewitt Foundation for Medical Research. This work was funded by the following grants to S.E.E. (R01AG049483); The Glenn Foundation for Medical Research Glenn Award for Research in Biological Mechanisms of Aging; a New Scholar in Aging Award from the Lawrence Ellison Foundation; and The Baxter Family Foundation; and an NIH grant to M.H. (R01 GM117466).

## AUTHOR CONTRIBUTIONS

Conceptualization, J.N.T., S.L.S., and M.H.; Methodology, J.N.T., S.L.S., R.C., S.L.B., S.E.E., and M.H.; Validation, J.N.T., S.L.S., R.C., S.L.B., S.E.E., and M.H.; Investigation, J.N.T., S.L.S., R.C., and S.L.B.; Writing – Original Draft, J.N.T., S.L.S., R.C., S.L.B., S.E.E., and M.H.; Writing – Review & Editing, J.N.T., S.L.S., R.C., S.L.B., S.E.E., and M.H.; Supervision, S.E.E. and M.H.; Funding Acquisition, J.N.T., S.E.E., and M.H.

## DECLARATION OF INTERESTS

The authors declare no competing interests.

## METHODS

### Mammalian Cell Culture

See Reagents Table for details of all cells and reagents employed. HEK 293T cells, HeLa cells, and N2A neuroblastoma cells were purchased from ATCC. Immortalized wild-type (WT) and STK4-deficient mouse embryonic fibroblasts (MEFs) were produced as described [6]. Cell lines were cultured in DMEM (Corning) supplemented with 10% fetal bovine serum (FBS, Gibco), referred to as “normal medium,” and were routinely checked for mycoplasma using the MycoScope PCR detection kit according to the manufacturer’s instructions (GenLantis).

Mouse hippocampal neurons were isolated from newborn BALB/c mice and cultured as described [26]. Briefly, hippocampi were dissected from 1- or 2-day-old mice, treated with papain (Worthington) for 15 min, and disrupted by aspirating through a micropipette tip 7 to 10 times. Dissociated neurons were resuspended in normal medium and plated in 24-well plates containing 12-mm glass coverslips pretreated with 50 µg/ml poly-L-lysine (Sigma) in borate buffer. The medium was exchanged for Neurobasal-A medium (Gibco) containing 2% B-27 (Gibco) and 0.25% GlutaMAX (Gibco) 1 h after plating.

### Plasmids and oligonucleotides

Plasmids encoding EGFP-tagged WT LC3B and phospho-mutants (T50A and T50E) were obtained as described [6]. Plasmids encoding HA-tagged LC3B proteins were generated by replacing the EGFP sequence in the EGFP plasmids with an HA-tag followed by a Tobacco etch virus protease sequence recognition site and a Flag-Tag (Genewiz) using *Age*I and *Hind*III (New England Biolabs). Lipidation-deficient LC3B proteins were generated by mutagenic PCR using Q5 Hot Start High-Fidelity Master Mix (New England Biolabs) to introduce the GGG to GCT change that leads to a glycine to alanine (G120A) mutation. Bacterial expression plasmids encoding His-tagged WT, T50A, and T50E LC3B proteins were produced as described [6]. A plasmid encoding mCherry-tagged FYCO1 was generated by cloning the mCherry-FYCO1 gene from a pBABE-puro-mCherry-FYCO1 plasmid (University of Dundee) into the plasmid backbone of EGFP-LC3B (Addgene) by amplification of fragments using Q5 Hot Start High-Fidelity Master Mix and ligation using a Gibson Assembly Master Mix (New England Biolabs). pLKO.1-puro plasmids encoding scrambled shRNA and five FYCO1-targeted shRNAs were purchased from Sigma Aldrich. For details on oligonucleotides refer to (**Table S1**).

### Generation of HeLa LC3B Knockout Cells

*LC3B* gene knockout (KO) was performed using the CRISPR-CAS9 system according to a published protocol [32]. Briefly, HeLa cells were transfected with two plasmids encoding GFP-CAS9 (Addgene) and an sgRNA targeting the first or the fourth exon of the *LC3B* gene. After 24 h, GFP-positive cells were sorted by FACS and single cells were placed in 96-well plates. Clones were grown for 2–3 weeks and analyzed for gene deletion by PCR. Gene-edited clones were expanded in 24-well plates, and LC3B protein deficiency was confirmed by western blotting and immunofluorescence microscopy. LC3B-KO cells were selected, expanded, and frozen before use in experiments.

### Gene Silencing, Transient Transfection, and Cell Treatment

Cells were reverse transfected with 10 nM of non-silencing control or gene-targeting siRNAs (siNS, siFYCO1, or siSTK4; Dharmacon) using Opti-MEM and lipofectamine RNAiMAX according to the supplier’s recommendations (Life Technologies). After 48 h, cells were collected and used for experiments. Transient plasmid transfections were performed using Opti-MEM and Lipofectamine 2000 (Life Technologies), and cells were collected for experiments 24 h after transfection. For experiments with cells expressing siRNAs and plasmids, cells were transfected with siRNAs for 24 h and then transfected with the plasmid of interest for an additional 24 h before analysis.

For autophagy flux assays, cells were washed three times with normal medium, normal medium containing the late-autophagy blocking compound bafilomycin A_1_ (BafA, Sigma) at 50 nM, or starvation medium (Earle’s Balanced Salt Solution (Life Technologies) before incubation in the same media for the indicated times at 37°C.

### Affinity purification

HeLa LC3B-KO cells, 293T cells, MEFs, or N2A cells were transiently transfected with plasmids (6.6 μg/10^6^ cells) encoding HA-LC3B-WT or HA-T50A for 24 h, incubated with normal medium or starvation medium for 1 h, and then lysed with IP lysis buffer (0.5% NP-40, 150 mM NaCl, 50 mM Tris HCl, pH 7.5, protease inhibitors [cOmplete, Roche], and phosphatase inhibitors [PhosStop, Roche]). Cell lysates were clarified by serial centrifugation (600 ×*g* for 3 min and 9300 ×*g* for 10 min) and diluted 1.25-fold with lysis buffer. An aliquot of the lysate (5%) was reserved as input for western blotting and the remaining lysate was incubated with anti-HA magnetic beads (Pierce) overnight at 4°C. Beads were washed seven times with wash buffer (0.1% NP-40, 150 mM NaCl, 50 mM Tris HCl, pH 7.5) and three times with 50 mM Tris HCl, pH 7.5. Samples were then processed for mass spectrometry or western blot analysis as described below. For immunoprecipitation of mCherry-FYCO1, the same protocol was performed using anti-RFP magnetic beads (MBL Life Science).

### Mass Spectrometry

Mass spectrometry was performed at the Proteomics Core of Sanford Burnham Prebys Medical Discovery Institute. Proteins immunoprecipitated as described above were subjected to on-bead digestion as previously described [33]. The total peptide concentration was determined using a NanoDrop spectrophotometer (Thermo Fisher) and the samples were then analyzed by LC-MS/MS using a Proxeon EASY nanoLC system (Thermo Fisher Scientific) coupled to a Q-Exactive Plus mass spectrometer (Thermo Fisher Scientific). Peptides were separated using an analytical C18 Acclaim PepMap column (75 µm × 250 mm, 2 µm vesicles; Thermo Scientific) and a 180-min gradient at a flow rate of 300 µl/min: 1% to 5% B in 1 min, 5% to 20% B in 139 min, 20% to 35% B in 30 min, and 35% to 45% B in 10 min (A = formic acid 0.1%; B = 80% acetonitrile + 0.1% formic acid). Mass spectrometry settings were as described previously [33]. Mass spectra were analyzed with MaxQuant software version 1.5.5.1. Peptides were searched against the *Homo sapiens* Uniprot protein sequence database (downloaded July 2018) and GPM cRAP sequences (commonly known protein contaminants), as described [33]. Statistical analysis was performed using an in-house R script (version 3.5.1) and SAINTq.

### Western blot analysis

Reserved input lysate (20 μg of total protein) or immunoprecipitated materials from affinity purifications were analyzed by standard western blotting protocols. Briefly, proteins were separated in Novex 4–12% acrylamide Bis-Tris gels (NuPage) and transferred to PVDF membranes. Membranes were blocked in Tris-buffered saline containing 0.05% Tween-20 (TBST) and either 5% milk or 3% bovine serum albumin (BSA), according to the antibody manufacturer’s recommendations, and then incubated for 2 h at room temperature with primary antibodies diluted in 1% milk or 3% BSA in TBST with gentle rocking. Blots were washed three times (total 30 min) and incubated with horseradish peroxidase (HRP)-conjugated anti-mouse or anti-rabbit secondary antibodies (Cell Signaling Technologies) for 1 h at room temperature with gentle rocking. Blots were washed again and developed with Pierce ECL Western Blotting substrate (Base, SuperSignal West Pico/Femto, depending on protein levels) and visualized using a Bio-Rad ChemiDoc Imaging System or with HyBlot CL autoradiography films (Denville). Quantification of band intensity was carried out using Image Lab (Bio-Rad) or ImageJ (National Institutes of Health) software.

### In Vitro Phosphorylation of LC3B and Co-immunoprecipitation of FYCO1

Aliquots of 3 μg of GST-LC3B (Viva Bioscience) were incubated alone or with 250 ng of His-STK4 (Millipore) in *in vitro* kinase reaction buffer (10 mM MgCl_2_, 1 mM EGTA, 1 mM dithiothreitol, 0.5 μm ATP, 20 mM Tris, pH 7.5) for 1 h at 30°C. The *in vitro* kinase reaction mixtures were incubated with 350 μl clarified lysate of HeLa LC3B-KO cells (8×10^5^ cell equivalents, prepared as described above) and glutathione-sepharose beads (Bioworld) for 3 h at 4°C on a rotator. As a negative control, the same procedure was performed with a mock *in vitro* reaction buffer lacking GST-LC3B and His-STK4. Sepharose beads were washed seven times with wash buffer (0.1% NP-40, 150 mM NaCl, 50 mM Tris HCl, pH 7.5) and eluted by heating in Laemmli sample buffer at 95°C for 10 min. Samples were passed through a 0.45 μm cellulose acetate filter (Costar) and then subjected to western blot analysis.

### Biolayer Interferometry

Cultures (500 ml) of Rosetta(tm) 2(DE3) pLysS *E. coli* (Millipore) expressing human LC3B-WT, T50A, or T50E [6] were grown to an optical density of 0.4–0.6 at 600 nm, and recombinant protein expression was induced by addition of 0.5 mM isopropyl-β-D-thiogalactopyranoside (VWR) for 16–18 h at 25°C. Bacteria were harvested by centrifugation and lysed with a microfluidizer in 50 mM Tris-HCl (pH 8.0), 300 mM NaCl, 30 mM imidazole, and 2 mM β-mercaptoethanol (BME). The lysates were clarified by centrifugation, filtered, incubated with Ni-NTA beads for 1 h, and eluted in the same buffer containing 250 mM imidazole. The sample was then dialyzed overnight in Snakeskin dialysis tubing (3500 kDa pore size) in 50 mM Tris-HCl (pH 8.5), 100 mM NaCl, and 5 mM BME. The protein was concentrated using Amicon-3.5K filters and subjected to size exclusion chromatography on a Superdex 200 column (GE Healthcare). The concentration of eluted proteins was measured at 280 nm using a Nanodrop spectrophotometer and the proteins were verified by Coomassie Blue staining and western blotting. Purified His-LC3B-WT, T50A, or T50E proteins were diluted to 3 µg/ml in Dulbecco’s phosphate-buffered saline (PBS; Gibco) containing 0.1% BSA and 0.02% Tween-20 and immobilized on NTA capture sensors (FortéBio). Human FYCO1 peptide (amino acids 1265– 1298, containing the LIR and adjacent residues) [4] was synthesized by Biomatik and reconstituted at 2 mg/ml in DMSO. The binding reaction was performed using two-fold serial dilutions of FYCO1 peptide (0.5–20 µM). The association and dissociation phases were analyzed for 60 s and the curves were inverted to fit them to a 1:1 model. Affinity constants were calculated from the kinetic constant after normalization for the buffer control. All binding measurements were performed on an Octet Red instrument (FortéBio) and the results were processed using Octet software 10.0.1 (FortéBio).

### Structural Representation of the LC3B–FYCO1-LIR Complex

The crystal structure of a *Mus musculus* complex between LC3B and FYCO1-LIR peptide (PDB 5WRD) [34] was represented in cartoon form using Pymol software. Distance measurements between T50 in LC3B and D1235 and D1236 adjacent to FYCO1-LIR (homologous to amino acids D1276 and 1277 in *Homo sapiens* FYCO1) were performed with Coot (Crystallographic Object-Oriented Toolkit) software and were found to be consistent with hydrogen bonding distances. The *Mus musculus* FYCO1-LC3B complex structure was chosen for representation because coverage of the FYCO1 peptide sequence was greater than in the human structure.

### Fluorescence and Confocal Microscopy

Cells (2×10^5^) were seeded in 24-well plates containing 12-mm glass coverslips, transiently transfected with the appropriate plasmids, and treated with normal or starvation medium as described above. After treatment, cells were fixed with 4% paraformaldehyde in PBS for 20 min and processed for fluorescence microscopy using standard protocols (https://www.cellsignal.com). Briefly, cells were incubated with primary antibodies against FYCO1 or autophagosomal and lysosomal proteins (see Reagents table) diluted in PBS containing 10% FBS and 0.2% saponin for 2 h at room temperature. Cells were then washed three times with PBS and incubated with Alexa Fluor 488- or 568-labeled secondary antibodies (Thermo Fisher) diluted at 1:1000 in PBS/10% FBS/0.2% saponin for 1 h at room temperature. Cells were washed three times in PBS, incubated with 4′,6-diamidino-2-phenylindole (DAPI) to label nuclei, and mounted with Prolong® Gold Antifade Reagent. Imaging was performed using a Zeiss LSM 710 NLO confocal microscope. For each condition, 15 to 20 images were acquired in randomly selected fields with a PlanApchromat 63X/1.4 DIC oil objective lens. An average of 10 to 15 stacks of 0.3 μm thickness were acquired.

### Image Analysis

Autophagosomes were quantified using Imaris 9.3.0 (Bitplane, Zurich). Confocal images of individual cells were processed using the software “spots” feature with an estimated diameter of 0.5 μm and then corrected using the “different spot sizes: region growing” feature. Thresholds were set using the “Quality” feature and determined manually. The local contrast threshold for region-growing spots was then manually set and maintained. Punctae were counted manually in 2–3 cells per condition and compared with the Imaris software counts to verify the accuracy and precision of measurements. The “surface” feature was used to reconstruct the nucleus of each cell using DAPI staining. Following generation of spots and surface objects, distance transformation was performed between the outer nuclear surface and the LC3B-positive punctae. Distances for individual punctae were exported to Excel (Microsoft) and processed. The distance from the nucleus of each punctum were averaged for each cell and the collective average of all punctae from all cells were represented as a single data point. Additionally, all distances for each condition were pooled and represented as relative frequency distributions with a best-fit line (non-linear regression, Gaussian curve) for interpretation. The same procedures were performed for LAMP-1 (endolysosomal marker; estimated punctum diameter = 0.33 μm). Co-localization data for FYCO1 and LC3B were processed using Imaris software using two methods. In the first method (main figures), spots were generated for LC3B and FYCO1 punctae in the manner described above. Then the “co-localize spots” software feature was used to identify LC3B and FYCO1 punctae located within very close proximity (<0.5 μm). These co-localized events were normalized to the total number of LC3B punctae and the data are presented as the mean percentage of LC3B/FYCO1 co-localization events. In the second method (supplemental figures), FYCO1- or LC3B-labeled channels were provided appropriate co-localization thresholds in the software “co-loc” feature using the “Polygon” tool. The same channel thresholds were maintained for all cells, and the most intense fluorescent overlap was quantified by Imaris as voxels (i.e., overlap between volumetric pixels in either channel) as described [35]. The voxel value for each cell was averaged for each condition and used as a single data point for statistical analysis.

### Live-Cell Imaging and LC3B Particle Tracking in HeLa Cells

HeLa LC3B-KO cells (2×10^5^) were seeded in 35-mm glass-bottom dishes (MatTek) for 24 h and transfected with GFP-tagged LC3B expression plasmids as described above. The medium was then replaced and the cells were imaged 2 h later with a Zeiss LSM 880 Rear Port Laser Scanning Confocal and Airyscan FAST microscope equipped with a stage pre-heated to 37°C. Approximately 9 to 10 cells per condition were randomly selected for imaging using a PlanApchromat 63X/1.4 DIC oil objective lens with single photon lasers emitting at 488 nm. Each cell was imaged for 3 min (360 frames, 2 frames/s) and, on average, 8–10 slices were acquired through a total cell thickness of ∼2.0–2.5 μm to ensure the complete punctum was captured for particle tracking in four dimensions. Following raw image acquisition, the movies were deconvoluted and a maximum Z-projection was generated using Zen Processing software. The movie files were further processed prior to particle tracking using Imaris image processing functions. The background subtraction function was set at 0.33 μm and a Gaussian-smoothing operation was used to remove the cytosolic GFP signal and to sharpen the visible punctae to improve particle tracking. GFP-positive punctae were marked using the spots feature of Imaris, with an estimated diameter of 0.5 μm for an individual punctum. An appropriate threshold was set to label all punctae visualized and was maintained for all movies analyzed. For particle tracking, the autoregressive motion algorithm was selected with 5–10 μm of maximum “predicted distance” and a maximum “gap distance” between 2 and 4 μm. Initial tracks were generated by Imaris and then manually curated and corrected to ensure each particle was assigned to the appropriate track for the duration of the movie. A reference frame was added over the nucleus and the general tracking statistics were exported to Excel for full processing. The final displacement relative to the reference frame labeled nucleus was subtracted from the initial displacement and each individual track was designated either “anterograde” or “retrograde.” The percent deviation from equal directional transport (50% anterograde, 50% retrograde) was pooled for each cell and the mean was taken as a single data point for statistical analysis. To quantify directional switches in particle movement, the complete track and trajectories showed by every single particle were displayed and directional switches were arbitrarily determined and quantified by two independent people using visual inspection of individual tracks.

### LC3B Particle Transport in Mouse Hippocampal Neurons

Hippocampal neurons were plated on 12-mm glass coverslips in 24-well plates, cultured for 8 to 10 days, and then transfected using 2 µL of Lipofectamine 2000 (Life Technologies) and 1.6 µg of DNA per well with the GFP-LC3B expression plasmids with or without scrambled shRNA or a mixture of five FYCO1-targeting shRNAs (Sigma Aldrich). The efficiency of shRNAs targeting *Fyco1* in mouse cells were verified in N2A neuroblastoma cells (**Figure S4D**), a proxy for primary neurons in which LC3B phospho-mutants displayed autophagy phenotypes similar to other cell types (**Figure S4E-F**). At 48 h post-transfection, the mid-axon region of neurons was imaged using a Nikon Ti-E Perfect Focus inverted microscope equipped with a total internal reflection fluorescence (TIRF) setup, an Andor iXon + DU897 EM camera, and a 100X/1.49 NA oil objective. A 488 nm laser was used for detection of GFP. Lasers were positioned at an angle for pseudo-TIRF acquisition. Each neuron was imaged for 5 min (300 frames, 1 frame/s). Axons were distinguished from dendrites by morphology [26], and the polarity of axons was determined by identification of soma and axonal termini for each movie. Axonal transport analysis was performed using KymoAnalyzer, a freely available ImageJ-based macro [36]. Briefly, kymographs were generated from time-lapse movies and particle trajectories were manually assigned from the kymograph images. Transport parameters, including direction, velocity, run length, time in motion, and pauses, were automatically calculated by KymoAnalyzer. A detailed description of all the transport parameters used in this study can be found in [36].

### Statistical Analysis

Statistical analysis and graph generation was conducted using Prism 8.0 software (GraphPad). One-way and two-way ANOVA were performed to analyze single-grouped datasets (Figures 1B, 1F, 2B, 4B, S1D, S2E and S4A) and double-grouped datasets (Figures 2D, 3C, 3E, 3G, S1C, S3E, and S4E), respectively. For all data sets, Tukey’s multiple comparison test with a single pooled variance and Geisser–Greenhouse correction was performed. P values are summarized using the GraphPad reporting method: 0.1234 (ns), 0.0332 (*), 0.0021 (**), 0.0002 (***), and <0.0001 (****). For the analysis of LC3B transport in primary hippocampal neurons, most axonal transport parameters did not follow a normal distribution and we therefore used the non-parametric permutation t test (rndttest function of MATLAB [Mathworks]; Figure 4D, 4G, S4B and S4F for those instances. For Figures 4E and S4F, parameter differences were analyzed using both the Rank Sum test (MATLAB [Mathworks]) and the Kolmogorov–Smirnov test (RStudio) adjusting p-values for multiple comparisons using Bonferroni.

## REAGENTS USED IN THIS STUDY

| REAGENT or RESOURCE | SOURCE | IDENTIFIER |
| --- | --- | --- |
| **Antibodies** |  |  |
| Rabbit polyclonal anti-FYCO1 | Atlas Antibodies | HPA035526 |
| Rabbit monoclonal anti-LC3B (D11) XP | Cell Signaling Technologies (CST) | 3868S |
| Rabbit polyclonal anti-MST1 | GeneTex | GTX109294 |
| Rabbit polyclonal anti-LAMP1 | Abcam | ab24170 |
| Rabbit polyclonal anti-p-MST1(Thr183)/MST2(Thr180) | CST | 3681S |
| Rabbit polyclonal anti pLC3B (Thr50) | 21<sup>st</sup> Century Biochemicals | (Wilkinson et al. 2015) |
| Mouse monoclonal anti-mCherry (1C51) | Novus Biologicals | NBP1-96752 |
| Mouse monoclonal anti-β-actin (C4) | Millipore Sigma | MAB1501 |
| Mouse monoclonal anti-HA (6E2) | CST | 2367S |
| Mouse monoclonal anti-α-tubulin (DM1A) | CST | 3873P |
| Goat anti-Rabbit HRP | CST | 7074s |
| Goat anti-Mouse HRP | CST | 7076s |
| Goat anti-Rabbit Alexa Fluor 488 | Thermo Fisher | A11034 |
| Goat anti-Rabbit Alexa Fluor 568 | Thermo Fisher | A11011 |
| Goat anti-Mouse Alexa Fluor 488 | Thermo Fisher | A11001 |
| Goat anti-Mouse Alexa Fluor 568 | Thermo Fisher | A11004 |
| **Bacterial and Virus Strains** |  |  |
| DH5α competent *E. coli* | Thermo Fisher | 18265017 |
| One Shot Stbl3 *E. coli* | Thermo Fisher | C7373-03 |
| Rosetta™ 2(DE3)pLysS *E. coli* | Millipore Sigma | 71403 |
| **Chemicals, Peptides, and Recombinant Proteins** |  |  |
| Dulbecco's Modified Eagle Medium (DMEM) | Corning | 10-013-CV |
| Fetal bovine serum (FBS) | Thermo Fisher | 26140049 |
| DMEM (for primary neurons) | Thermo Fisher | 11965 |
| FBS (for primary neurons) | Thermo Fisher | 10082147 |
| Poly-L-lysine | Millipore Sigma | P5899 |
| Neurobasal-A medium | Thermo Fisher | 10888022 |
| B-27 | Thermo Fisher | 17504044 |
| GlutaMAX | Thermo Fisher | 35050061 |
| Earle's Balanced Salt Solution (EBSS): starvation medium | Thermo Fisher | 24010043 |
| Lipofectamine 2000 | Thermo Fisher | 11668019 |
| Opti-MEM | Thermo Fisher | 31985-070 |
| Bafilomycin A1 | Millipore Sigma | 88899-55-2 |
| DAPI | Millipore Sigma | D9542 |
| RNAiMAX | Thermo Fisher | 13778150 |
| Saponin, from Quillaja pract. | Acros Organics | 74499-23-3 |
| N3500 Nonidet-P40 substitute | US Biological | 9036-19-5 |
| Bovine serum albumin (BSA), protease free | Akron | AK1391-0100 |
| cOmplete Protease Inhibitor Cocktail tablets | Millipore Sigma | 11697498001 |
| PhosSTOP | Millipore Sigma | 4906845001 |
| Pierce ECL Western Blotting Substrate | Thermo Fisher | 32209 |
| SuperSignal™ West Pico PLUS Chemiluminescent Substrate | Thermo Fisher | 34577 |
| SuperSignal™ West Femto Maximum Sensitivity Substrate | Thermo Fisher | 34095 |
| Prolong® Gold Antifade Reagent | CST | 9071 |
| FYCO1 peptide (1265–1298 amino acids of the human protein) | Biomatik | N/A |
| GST-LC3B | Viva Bioscience | V32470-0500 |
| His-STK4 | Millipore Sigma | 14-624 |
| **Critical Commercial Assays** |  |  |
| Q5® Hot Start High-Fidelity 2X Master Mix | New England Biolabs | M0494S |
| Gibson Assembly® Master Mix | New England Biolabs | E2611L |
| MycoScope PCR Mycoplasma Detection Kit | Genlantis | MY01100 |
| **Experimental Models: Cell Lines** |  |  |
| Human: HEK 293T cells | ATCC | CRL-3216 |
| Human: HeLa cells | ATCC | CCL-2 |
| Human: HeLa cells lacking endogenous LC3B | This paper | N/A |
| Mouse: Immortalized MEFs | (Wilkinson et al. 2015) | N/A |
| Mouse: Immortalized STK3<sup>+/-</sup>4<sup>-/-</sup> MEFs | (Wilkinson et al. 2015) | N/A |
| Mouse: Primary hippocampal neurons | Encalada; BALB/c | N/A |
| Mouse: Neuro2a (N2A) neuroblastoma cells | ATCC | CCL-131 |
| **Oligonucleotides** |  |  |
| ON-TARGETplus Control Pool: Non-Targeting pool | Dharmacon | D-001810-10-05 |
| ON-TARGETplus FYCO1 siRNA | Dharmacon | LQ-014350-01-0002 |
| ON-TARGETplus SMARTpool: Human STK4 | Dharmacon | L-004157-00-0005 |
| **Recombinant DNA** |  |  |
| Plasmid: pHA-tag | This paper | N/A |
| Plasmid: pHA-LC3B | This paper | N/A |
| Plasmid: pHA-LC3B-T50A | This paper | N/A |
| Plasmid: pHA-LC3B-T50E | This paper | N/A |
| Plasmid: pHA-LC3B-G120A | This paper | N/A |
| Plasmid: pHA-LC3B-T50A-G120A | This paper | N/A |
| Plasmid: pHA-LC3B-T50E-G120A | This paper | N/A |
| Plasmid: pHA-STK4 | This paper | N/A |
| Plasmid: pEGFP-LC3B | This paper | N/A |
| Plasmid: pEGFP-LC3B-T50A | Wilkinson 2015 | N/A |
| Plasmid: pEGFP-LC3B-T50E | Wilkinson 2015 | N/A |
| Plasmid: pBABE-mCherry-FYCO1 | Univ. of Dundee | DU45204 |
| Plasmid: pSpCas9(bb)-2A-GFP | Addgene | 48138 |
| Plasmid: pSpCas9(bb)-2A-GFP-LC3B Human A | This paper | N/A |
| Plasmid: pSpCas9(bb)-2A-GFP-LC3B Human B1 | This paper | N/A |
| Plasmid: pSpCas9(bb)-2A-GFP-LC3B Human B2 | This paper | N/A |
| pLKO.1-puro non-targeting (scramble) shRNA Control | Millipore Sigma | SHC016-1EA |
| pLKO.1-puro FYCO1 shRNA 1 | Millipore Sigma | TRCN0000184601 |
| pLKO.1-puro FYCO1 shRNA 2 | Millipore Sigma | TRCN0000183567 |
| pLKO.1-puro FYCO1 shRNA 3 | Millipore Sigma | TRCN0000179990 |
| pLKO.1-puro FYCO1 shRNA 4 | Millipore Sigma | TRCN0000178746 |
| pLKO.1-puro FYCO1 shRNA 5 | Millipore Sigma | TRCN0000183006 |
| **Software and Algorithms** |  |  |
| SAINTq software | SourceForge | N/A |
| In house R script (ver. 3.5.1, 64-bit) | SBP: Proteomics | N/A |
| Octet software 10.0.1 | FortéBio | N/A |
| PyMOL 2.3 | Schrödinger | N/A |
| Imaris 9.3.0 | Bitplane | N/A |
| ImageJ | NIH | N/A |
| Zen lite processing software | Zeiss | N/A |
| Prism 8 | GraphPad | N/A |
| Microsoft Office: Excel, PowerPoint, Word | Microsoft | N/A |
| RStudio | RStudio Inc | N/A |
| Image Lab | Thermo Fisher | N/A |
| **Other Reagents** |  |  |
| Pierce Anti-HA magnetic beads | Thermo Fisher | 88836 |
| Anti-RFP magnetic beads | MBL Life Science | M165-11 |
| 1-1/2 Micro glass coverslips – 12 mm diameter | Electron Microscopy Sciences | 72230-01 |
| Premium microscope slides (Superfrost) | Thermo Fisher | 12-544-7 |
| Spin-X centrifuge tube filter 0.45 µm cellulose acetate | Corning Inc. | 8162 |
| NuPAGE 4–12% Bis-Tris gels 1 mm × 10 wells | Thermo Fisher | NP0321BOX |
| Premium autoradiography film (5 × 7 in.) | Denville Scientific | E3012 |
| Immobilon-P membrane, PVDF, 0.45 µm, 8.5 cm × 10 m roll | Millipore Sigma | IPVH85R |

## SUPPLEMENTAL INFORMATION

**Figure S1.**
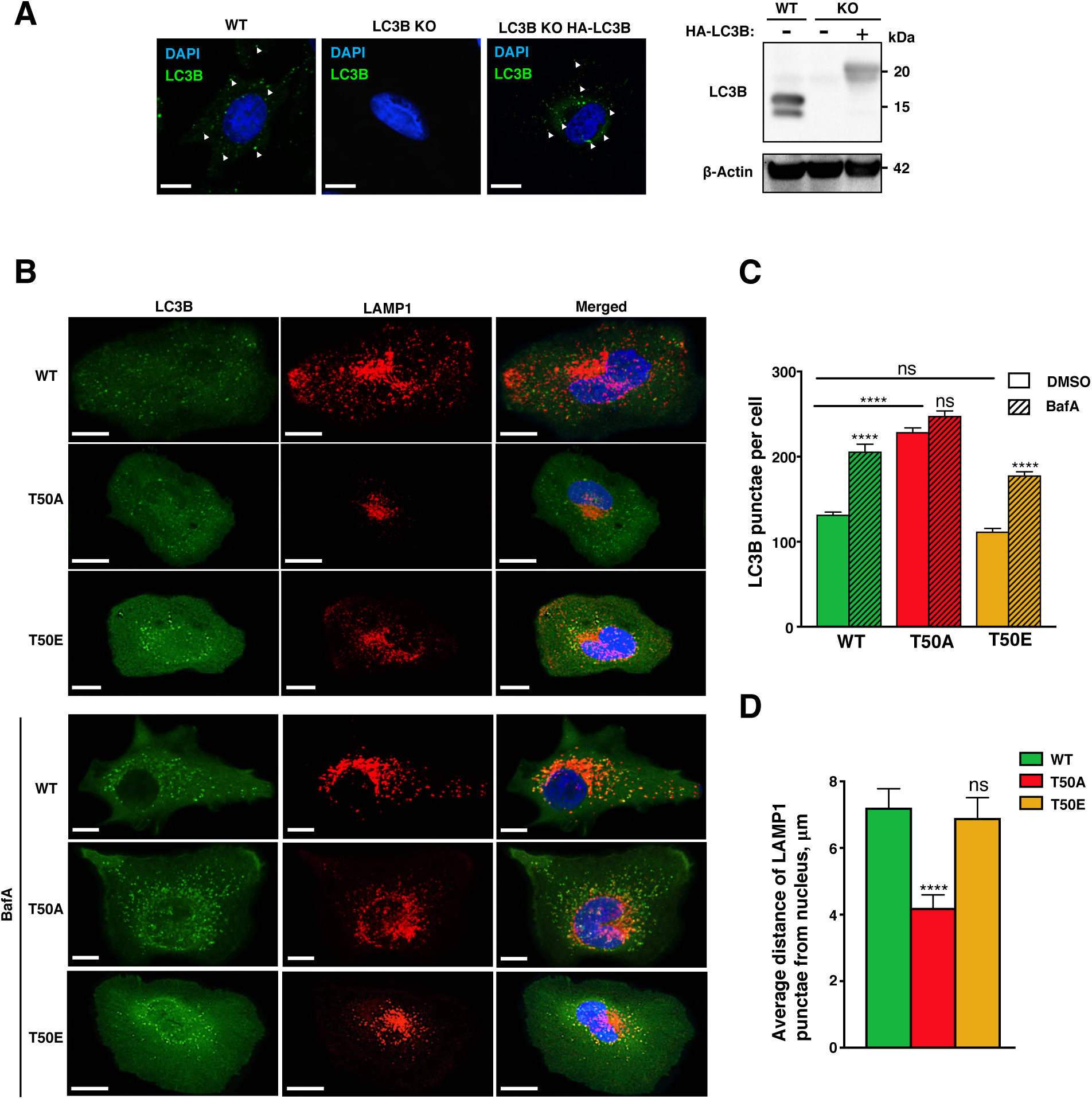
Influence of LC3B phosphorylation on autophagy flux and lysosomal positioning in HeLa LC3B-KO cells. (**A**) Representative immunofluorescence micrographs of HeLa WT, LC3B-KO, or LC3B-KO transfected with HA-LC3B. Cells were stained with anti-HA antibody (green). Arrowheads indicate LC3B-positive punctae (potential autophagosomes). Right panel shows western blot of LC3B in the WT and LC3B-KO cells (n=2). (**B**) Representative immunofluorescence micrographs of LC3B-KO HeLa cells transiently expressing HA-LC3B-WT, HA-T50A, or HA-T50E and incubated in normal medium (upper panels) or medium containing 50 nM bafilomycin A1 (BafA; lower panels) for 6 h. Cells were stained with anti-HA (green) and anti-LAMP1 (red) antibodies. Nuclei were stained with DAPI (blue). (**C**) Quantification of LC3B punctae in images shown in (B). Mean ± SEM of n=15 cells from two experiments. ****p<0.00005 by two-way ANOVA. (**D**) Quantification of the distance between lysosomes and the cell nucleus. Mean ± SEM of n=8–18 cells from two experiments. ***p<0.0005 by two-way ANOVA.

**Figure S2.**
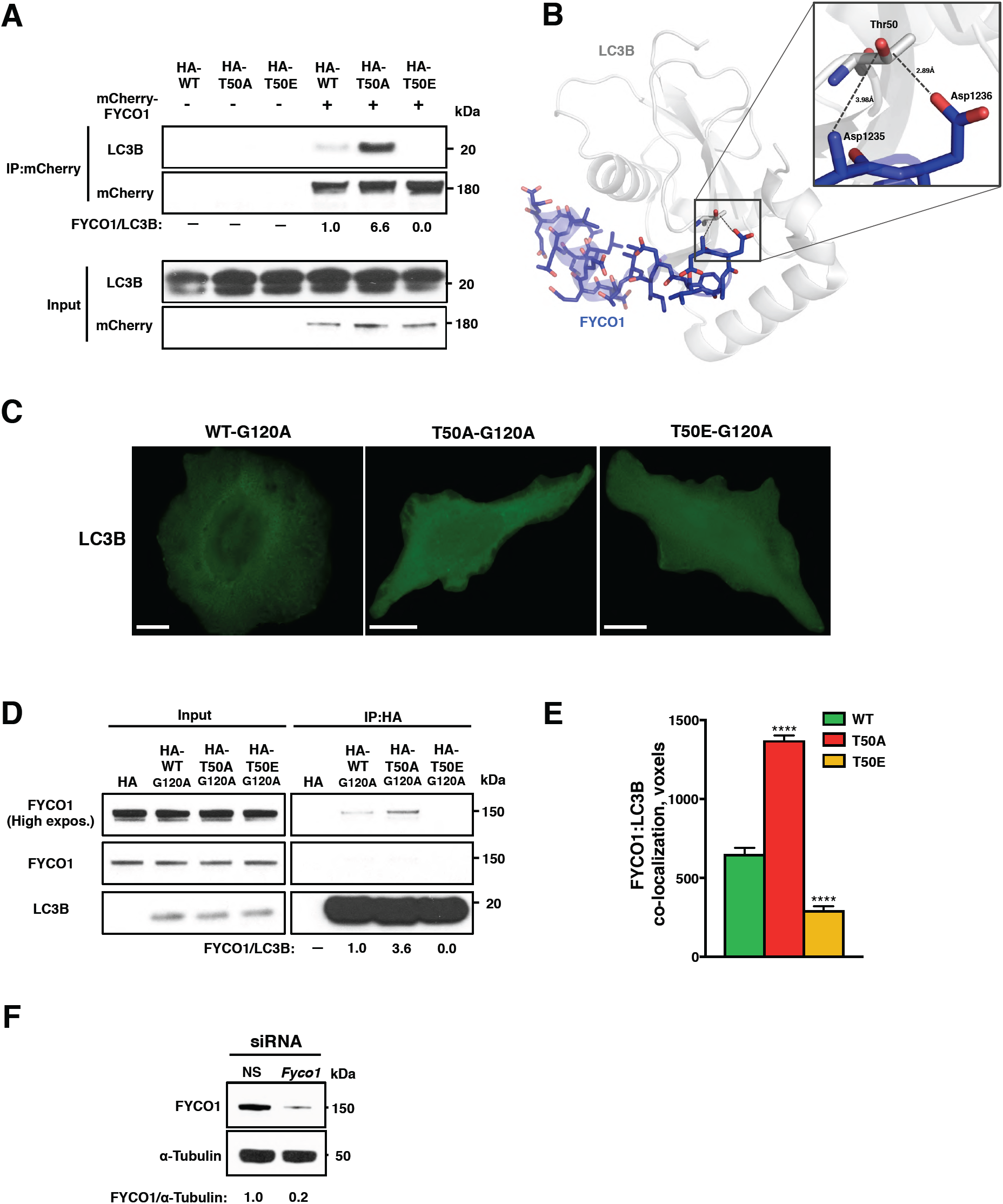
LC3B phosphorylation modulates FYCO1–LC3B interactions in an LC3B lipidation-independent manner. (**A**) Representative western blot of LC3B co-immunoprecipitated with mCherry-FYCO1 in HeLa LC3B-KO cells expressing the indicated HA-tagged LC3B proteins. FYCO1/LC3B values indicate the ratio of proteins (n=2). (**B**) Crystal structure of LC3B protein (gray) complexed with FYCO1-LIR (blue) (PDB 5WRD, *Mus musculus*) [34]. Potential hydrogen bonds (black dotted lines) between LC3B threonine 50 and two aspartic acid residues adjacent to the FYCO1-LIR are shown. (**C**) Representative immunofluorescence images of HeLa LC3B-KO cells expressing lipidation-deficient (G120A) HA-tagged LC3B proteins. Scale bar = 10 μm. Punctae counts: WT = 0.4 ± 0.2, T50A = 0.7 ± 0.2, T50E = 0.2 ± 0.1. n=10 cells from two experiments. (**D**) Representative western blot of FYCO1 co-immunoprecipitated with HA-tagged LC3B constructs (WT, T50A, T50E) containing the G120A mutation, which prevents lipidation, in HeLa LC3B-KO cells. The mutant proteins were expressed at levels similar to those of their lipidation-proficient counterparts. FYCO1/LC3B values indicate the ratio of proteins (n=2). (**E**) Co-localization between FYCO1 and LC3B punctae “voxels” (volumetric pixels) in HeLa LC3B-KO cells expressing LC3B-WT, T50A, or T50E proteins. Mean ± SEM of n=13–26 cells from three experiments. ***p<0.0002 by two-way ANOVA. (**F**) Representative western blot of FYCO1 in HeLa cells expressing control (NS) or FYCO1-targeting siRNAs. FYCO1/α-tubulin values indicate the ratio of proteins (n=2).

**Figure S3.**
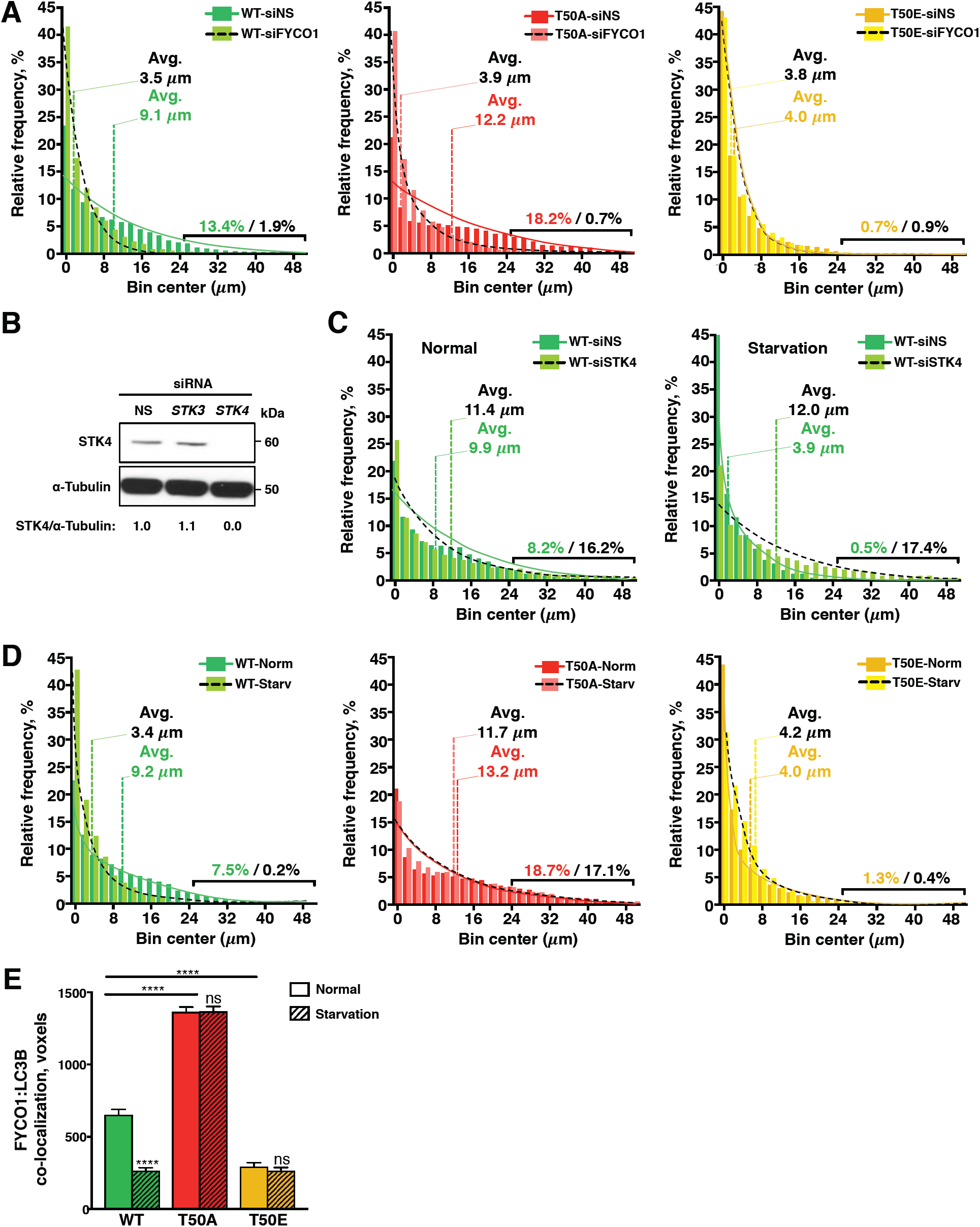
Effect of starvation and STK4 silencing on autophagosome positioning and FYCO1–LC3B colocalization. (**A**) Relative frequency distribution of the distance between autophagosomes and the nucleus in HeLa cells co-expressing LC3B proteins and control (siNS) or FYCO1-targeting siRNA. The mean distances are shown and a non-linear regression best-fit line was overlaid on each distribution. The percentage of LC3B punctae located at distances greater than 25 µm from the nucleus is also shown. n=24–34 cells from three experiments. (**B**) Representative western blot of STK4 levels in HeLa LC3B-KO cells treated with control (siNS), *Stk3-*targeting, or *Stk4-*targeting siRNAs. STK4/α-tubulin values indicate the ratio of proteins (n=2). (**C**) As for (A) except HeLa LC3B-KO cells were transfected with control or STK4-targeting siRNA. n=24–34 cells from three experiments. (**D**) As for (A) except HeLa cells were incubated in normal or starvation medium. n=24–34 cells from three experiments. (**E**) FYCO1-LC3B co-localization (“voxels”) in HeLa LC3B-KO cells expressing the indicated LC3B proteins after incubation in normal or starvation medium. Mean ± SEM of 13–27 cells from two experiments. ***p<0.0002 by two-way ANOVA.

**Figure S4.**
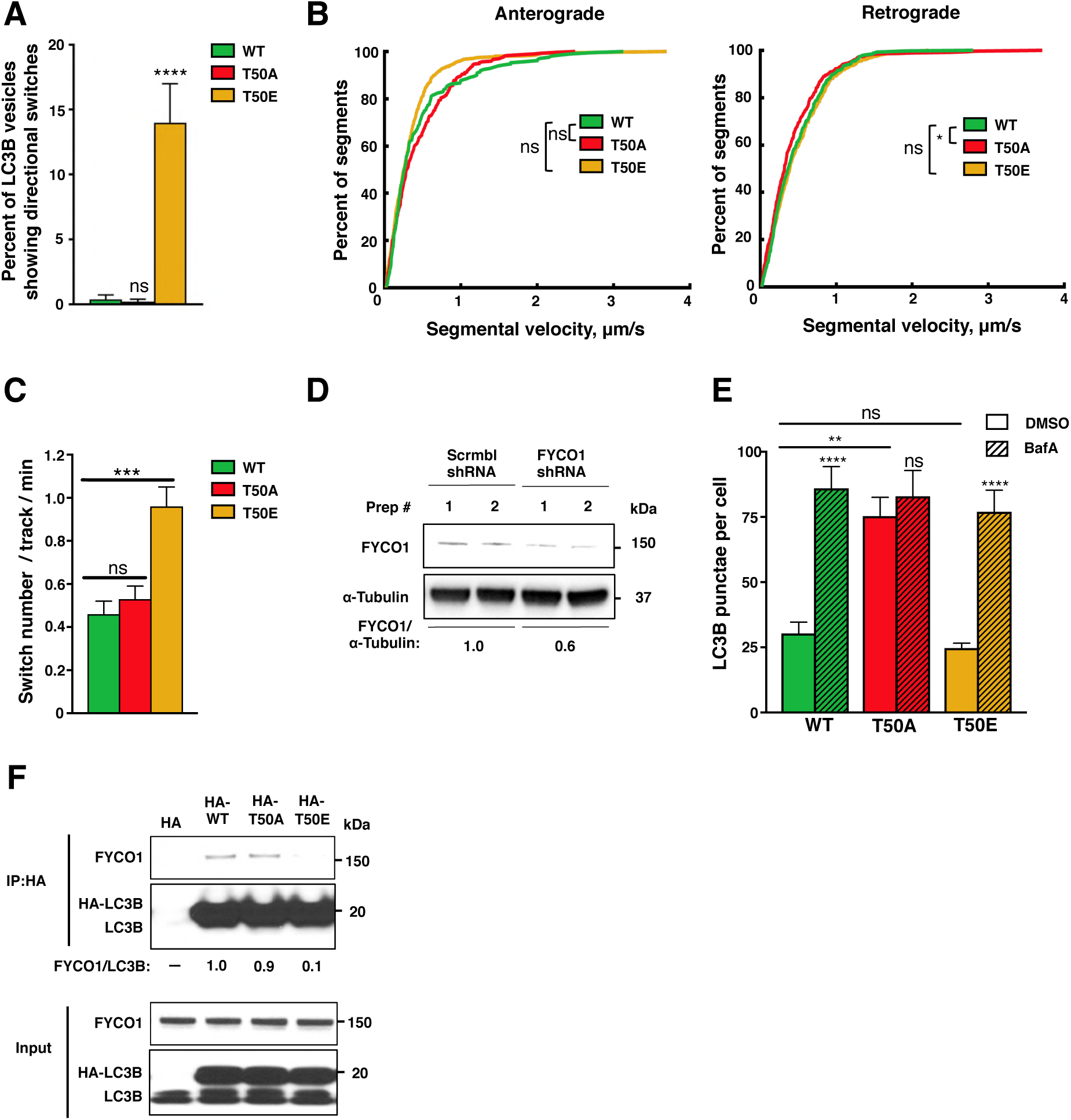
Conserved effects of LC3B phosphorylation status on autophagy and autophagosome transport in neuronal cells. (**A**) Average percentage of LC3B-positive vesicles showing directional switches in HeLa LC3B KO cells. Mean ± SEM of n=9–10 cells from two experiments. ****p<0.0001 by one-way ANOVA. (**B**) Cumulative frequency graphs of anterograde and retrograde segmental velocities of GFP-LC3B-positive vesicles in mouse primary neurons. n=26–29 cells from two experiments. *p<0.05 by Rank Sum test. (**C**) Mean directional switches per particle per min of GFP-LC3B-positive vesicles in primary mouse neurons. Mean ± SEM of n=26–29 cells from two experiments. *p<0.05 by non-parametric permutation Student’s t-test. (**D**) Representative western blot of FYCO1 levels in mouse N2A neuroblastoma cells expressing control (Scrmbl) or *Fyco1*-targeting shRNAs. FYCO1/α-tubulin values indicate the ratio of proteins (n=2). (**E**) Quantification of LC3B punctae in N2A neuroblastoma cells transiently expressing HA-tagged LC3B proteins. Cells were incubated for 6 h in medium containing vehicle (DMSO) or 50 nM bafilomycin A1 (BafA). Mean ± SEM of 9–12 cells from two experiments. ****p<0.00005 by two-way ANOVA (**F**) Representative western blot of FYCO1 co-immunoprecipitated with HA-tagged LC3B proteins in N2A cells. FYCO1/LC3B values indicate the ratio of proteins (n=3).

**Video S1, S2, and S3. Directional transport of LC3B-WT, T50A, and T50E vesicles in HeLa LC3B-KO cells.**

**Video S4, S5, and S6. Directional transport of LC3B-WT, T50A, and T50E vesicles in mouse primary neurons.**

**Table S1.**
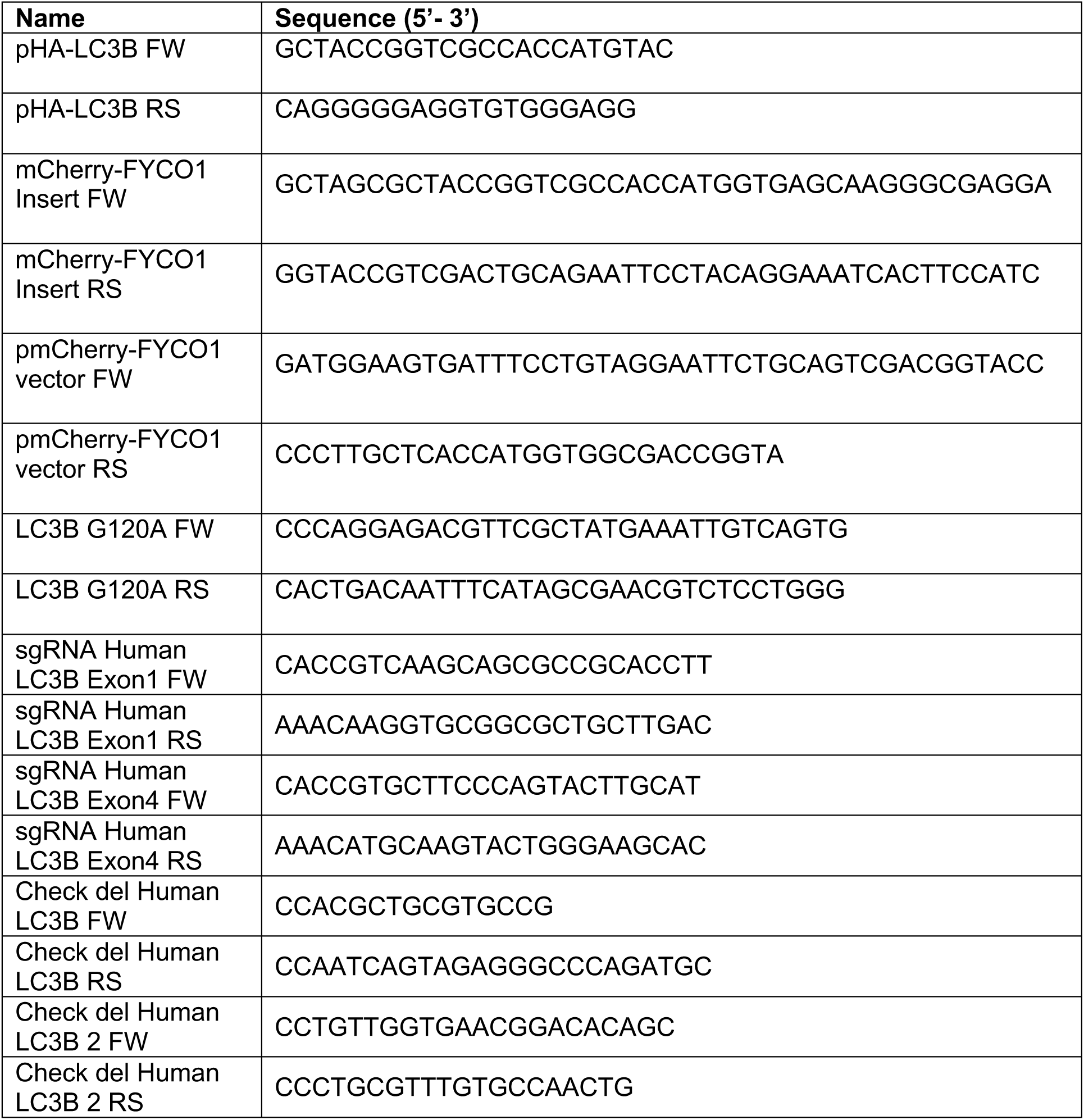
Oligonucleotides Used in This Study, Related to Methods.

## Notes

### Competing Interest Statement

The authors have declared no competing interest.

